# Epigenetic Age Acceleration in Surviving versus Deceased COVID-19 Patients with Acute Respiratory Distress Syndrome following Hospitalization

**DOI:** 10.1101/2023.07.18.549478

**Authors:** Yosra Bejaoui, Fathima Humaira Amanullah, Mohamad Saad, Sarah Taleb, Martina Bradic, Andre Megarbane, Ali Ait Hssain, Charbel Abi Khalil, Nady El Hajj

**Author notes:** **Correspondence:** Charbel Abi Khalil, Department of Genetic Medicine, Weill Cornell Medicine-Qatar, Doha-Qatar, and Nady El Hajj, College of Health and Life Sciences, Hamad Bin Khalifa University, Qatar Foundation, Education City, Doha, Qatar. Equal collaboration (Charbel Abi Khalil, Nady El Hajj).

## Abstract

Aging has been reported as a major risk factor for severe symptoms and higher mortality rates in COVID-19 patients. Molecular hallmarks such as epigenetic alterations and telomere attenuation reflect the biological process of aging. Epigenetic clocks have been shown to be valuable tools for measuring biological age in a variety of tissues and samples. As such, these epigenetic clocks can determine accelerated biological aging and time-to-mortality across various tissues. Previous reports have shown accelerated biological aging and telomere attrition acceleration following SARS-CoV-2 infection. However, the effect of accelerated epigenetic aging on outcome (death/recovery) in COVID-19 patients with Acute Respiratory Distress Syndrome (ARDS) has not been well investigated. In this study, we measured DNA methylation age and telomere attrition in 87 severe COVID-19 cases with ARDS under mechanical ventilation. Furthermore, we compared dynamic changes in epigenetic aging across multiples time-points until recovery or death. Epigenetic age was measured using the Horvath, Hannum, DNAm skin and blood, GrimAge, and PhenoAge clocks, whereas telomere length was calculated using the surrogate marker DNAmTL. Our analysis revealed significant accelerated epigenetic aging but no telomere attrition acceleration in severe COVID-19 cases. In addition, we observed epigenetic age deceleration at inclusion vs end of follow-up in recovered but not in deceased COVID-19 cases using certain clocks. When comparing dynamic changes in epigenetic age acceleration (EAA), we detected higher EAA using both the Horvath and PhenoAge clocks in deceased vs recovered patients. The DNAmTL measurements revealed telomere attrition acceleration in deceased COVID19 patients between inclusion and end of follow-up as well as a significant change in dynamic telomere attrition acceleration when comparing patients who recovered vs those who died. In conclusion, EAA and telomere attrition acceleration was associated with treatment outcome in hospitalized COVID-19 Patients with ARDS. A better understanding of the long-term effects of EAA in COVID19 patients and how they might contribute to Long COVID symptoms in recovered individuals is urgently needed.

## Background

The global outbreak of COVID-19 resulted in a significant public health crisis with wide-ranging implications. According to the world health organization, over 767 million confirmed cases and 7 million deaths have been attributed to COVID-19 as of July 2023 (https://covid19.who.int). COVID-19 is caused by an enveloped single-stranded positive RNA virus known as severe acute respiratory coronavirus 2 (SARS-CoV-2), which first emerged in Wuhan City, China, in late 2019 (1). The main causes of death in patients with COVID-19 are respiratory failure and multiorgan dysfunction due to impaired immune response and uncontrolled inflammatory processes (2). Patients with severe types of COVID-19 frequently experience respiratory failure and may often require mechanical ventilation (3). Although the major signs and symptoms of COVID-19 are currently well known, there is increasing evidence that the virus may have long-term effects and may impact many different aspects of human health. Those ongoing health problems following initial COVID-19 infection are commonly referred to as Long COVID or Post-COVID Conditions (4).

Chronological age is one of the well-established prognostic factors in patients with COVID-19 independent of other age-related comorbidities such as diabetes and cardiovascular diseases (5). COVID-19 poses a disproportionate threat to older adults due to the increased risk of disease severity, mortality rates, and long-term consequences (6) in contrast to infants and children who often exhibit milder clinical symptoms. Aging is a complex biological process characterized by a progressive decline in physiological function and increased disease susceptibility. Aging is defined by specific hallmarks such as genomic instability, loss of proteases, stem cell exhaustion, telomere attrition, and epigenetic alterations (7). The most studied type of epigenetic alteration is DNA methylation, which is an addition of a methyl group to the 5^th^ carbon position of the cytosine ring to form 5-methylcytosine. This modification mainly occurs within the context of a CpG dinucleotide and is known to regulate gene expression (8). DNA methylation signatures are known to be impacted by environmental exposures and are strongly correlated with aging in multiple tissues (9-11).

DNAm age often referred to as epigenetic age is a measure of biological age based on DNA methylation of specific CpG sites that reflect environmental exposures and disease risks (12). Epigenetic age acceleration has been reported to be associated with multiple diseases including cancer, diabetes, Alzheimer’s disease, HIV infection, and certain progeroid syndromes (10, 12-15). Several epigenetic clocks have been developed to estimate epigenetic age (16), such as the pan-tissue Horvath clock based on 353 CpGs (17), the PhenoAge clock based on 513 CpGs (18), and the GrimAge based on 1030 CpGs associated with physiological and stress risk factors (19). In addition, DNA methylation biomarkers can be used to estimate telomere length, which is another hallmark of aging. The DNAm telomere length estimator (DNAmTL) is based on methylation measurement of 140 CpG sites (20). Recent research has shown a potential link between COVID-19 and accelerated aging, as demonstrated by changes in the epigenetic age (21). Given the severity and global impact of the COVID-19 pandemic, investigating the potential influence of SARS-CoV-2 infection on accelerated biological aging is of significant public health relevance. Studies have reported intriguing findings, with indications that COVID-19 patients exhibit accelerated epigenetic aging compared to healthy individuals of similar chronological age (21). Furthermore, a significant acceleration of telomere attrition was observed comparing healthy vs COVID-19 patients. Severe COVID-19 infection often leads to respiratory failure requiring prolonged mechanical ventilation. A recent study by Cao *et al*. reported accelerated epigenetic aging to be associated with the severity of COVID-19; however, the underlying mechanisms and the broader implications of these observations remain poorly understood.

This study aimed to comprehensively examine epigenetic age acceleration in COVID-19 patients with acute respiratory distress syndrome (ARDS) and how it relates to outcome (survival or death) following hospitalization. In addition, we studied dynamic epigenetic aging across severe COVID-19 disease phases starting from intensive care unit (ICU) admission until death or recovery. The epigenetic age acceleration was calculated using several clocks including Horvath, Hannum, DNAm skin and blood, GrimAge, and PhenoAge clocks. Furthermore, telomere length was estimated via the surrogate marker DNAmTL to elucidate the impact of COVID-19 on telomere attrition.

## Results

### Epigenetic age acceleration in COVID-19 patients at baseline

We measured epigenetic age using five different epigenetic clocks in whole blood samples collected from 87 hospitalized COVID-19 patients with ARDS under mechanical ventilation at inclusion time-point T1 and 21 control samples. A strong positive correlation was observed when comparing epigenetic age vs chronological age using the different clocks including Horvath (r=0.87, p=2.5e-34), SkinBlood (r=0.92, p=6.3e-45), Hannum (r=0.88, p=4.7e-36), PhenoAge (r=0.83, p=1.2e-28), and GrimAge (r=0.93, p=7e-48) (Supp. Fig 1). Our analysis on Epigenetic Age Acceleration (EAA) in COVID-19 patients revealed significant acceleration when compared to control samples in three different epigenetic clocks: Hannum clock (p=0.0168), PhenoAge clock (p=0.0048), and GrimAge clock (p=0.002) after adjusting for BMI and gender (Fig1).

**Fig. 1.**
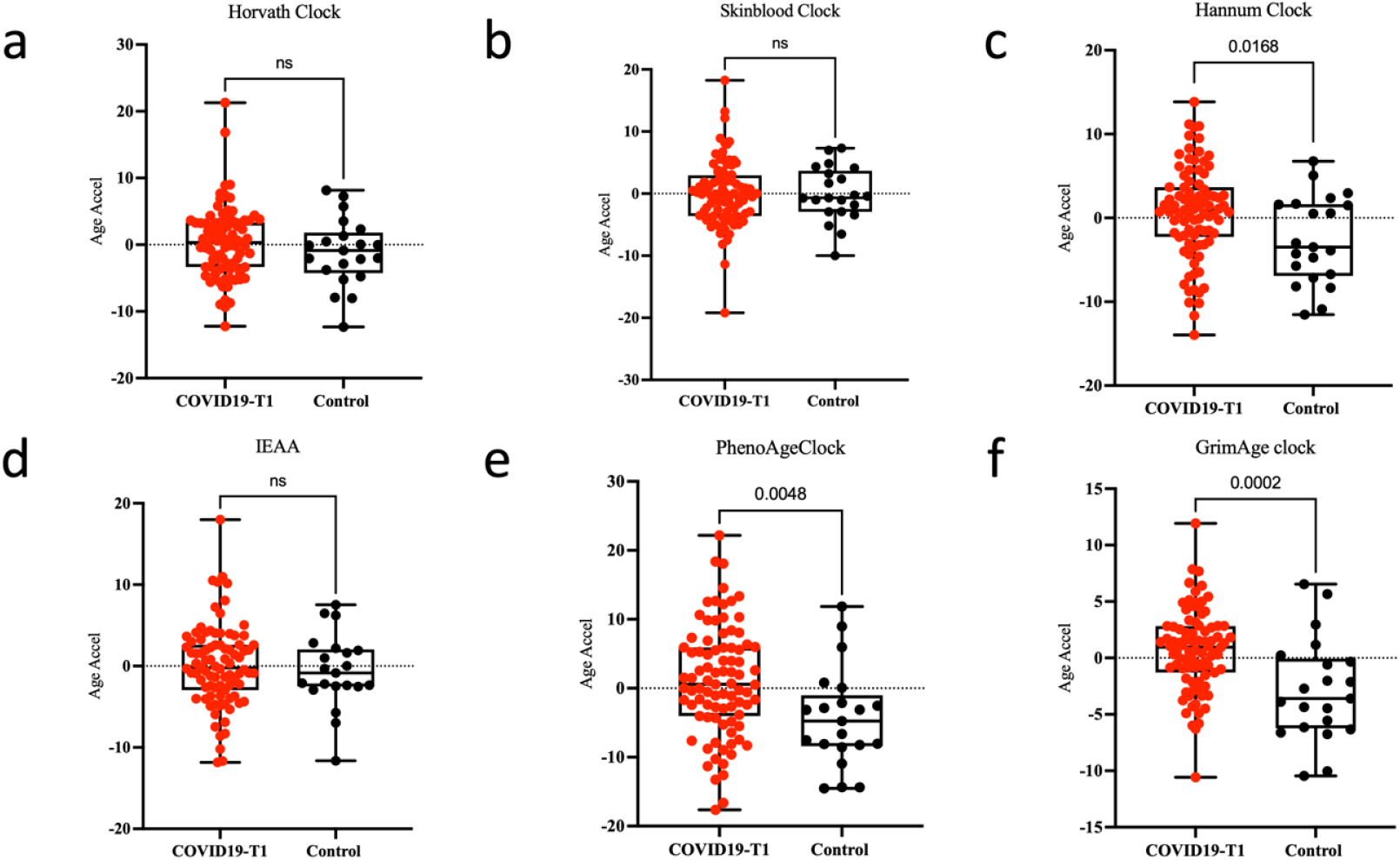
Accelerated epigenetic aging in COVID-19 patients T1 vs controls. DNAm age acceleration in six epigenetic clocks (a-f) in the peripheral blood from 21 control samples and 87 severe COVID-19-T1patients. The y-axis shows the epigenetic age. The p-value for each t-test is shown above the corresponding line. In the box plots, the lower and upper hinges indicate the 25th and 75th percentiles and the black line within the box marks the median value. Ns: non-significant.

### Epigenetic age acceleration in deceased and recovered critically ill COVID-19 patients

In total, 78% of the COVID-19 patients survived and were discharged from ICU at different time points. Patients who died were on average older compared to those who recovered (p <L0.001) (**Table 1**). In addition, they had a lower BMI, and were more likely to have hypertension and chronic kidney disease, yet those difference were not significant between both groups (p>0.05). First, we assessed epigenetic age acceleration in the subgroup of COVID-19 patients who survived following admission to the ICU. Here, we examined DNAmAge at inclusion (T1) for 50 COVID-19 surviving patients to their last recorded DNAmAge before recovery. Among the different epigenetic clocks used in our study, the Horvath clock (p=0.0017), Hannum clock (p<0.0001) and PhenoAge clock (p=0.009) revealed a significant decrease in epigenetic age acceleration at the last recorded time prior to recovery (Fig2). Next, we measured epigenetic age acceleration in the subset of 14 COVID-19 patients who died following ICU admission. Comparing DNAmAge at inclusion to the last recorded DNAmAge before death revealed no significant epigenetic age acceleration using the different epigenetic clocks (Fig. 3). We additionally compared DNAmAge in COVID-19 patients who died to those who recovered at both baseline level and the final timepoint of follow-up. This analysis revealed no epigenetic age acceleration using different epigenetic clocks in both comparisons (Fig. 4).

**Table 1.** Clinical characteristics of surviving and deceased COVID-19 patients following hospitalization. Data are represented as numbers (%) per each category for categorical variables and as median and 1st and 3rd quartile for continuous variables. ICU= Intensive Care Unit, LoS= Length of Stay, MV: Mechanical Ventilation.

|  | COVID-19 Survived (N=68) | COVID-19 Died (N=19) | Controls (N=21) |
| --- | --- | --- | --- |
| **Age** | 47 (41-53.5) | 58 (52.5-63.5) | 40 (34.5-47) |
| **Gender (male)** | 100% | 93% | 100% |
| **BMI (kg/m2)** | 27.44 (25.33-31.39) | 25.7 (24.28-29.64) | 28.7 (26-30.1) |
| **Duration of MV (days)** | 7 (4-13.5) | 20 (14.5-26.5) | - |
| **ICU LoS (days)** | 14 (9.75-26.25) | 24 (15-30.5) | - |
| **Hospital LoS (days)** | 30 (22-46) | 25 (17-35) | - |
| **Nosocomial infections** | 10% | 21.05% | - |
| **Convalescent plasma therapy** | 51% | 79% | - |
| **Diabetes status** |  |  |  |
| Non diabetes | 50% | 63% | - |
| Pre-diabetes | 2% | 5% | - |
| Diabetes | 47% | 32% | - |
| **Hypertension** | 43% | 47% | - |
| **Coronary artery disease** | 5.9% | 5.3% | - |
| **Chronic kidney failure** | 4.4% | 11% | - |
| **Chronic heart failure** | 1.5% | 0% | - |

**Fig. 2.**
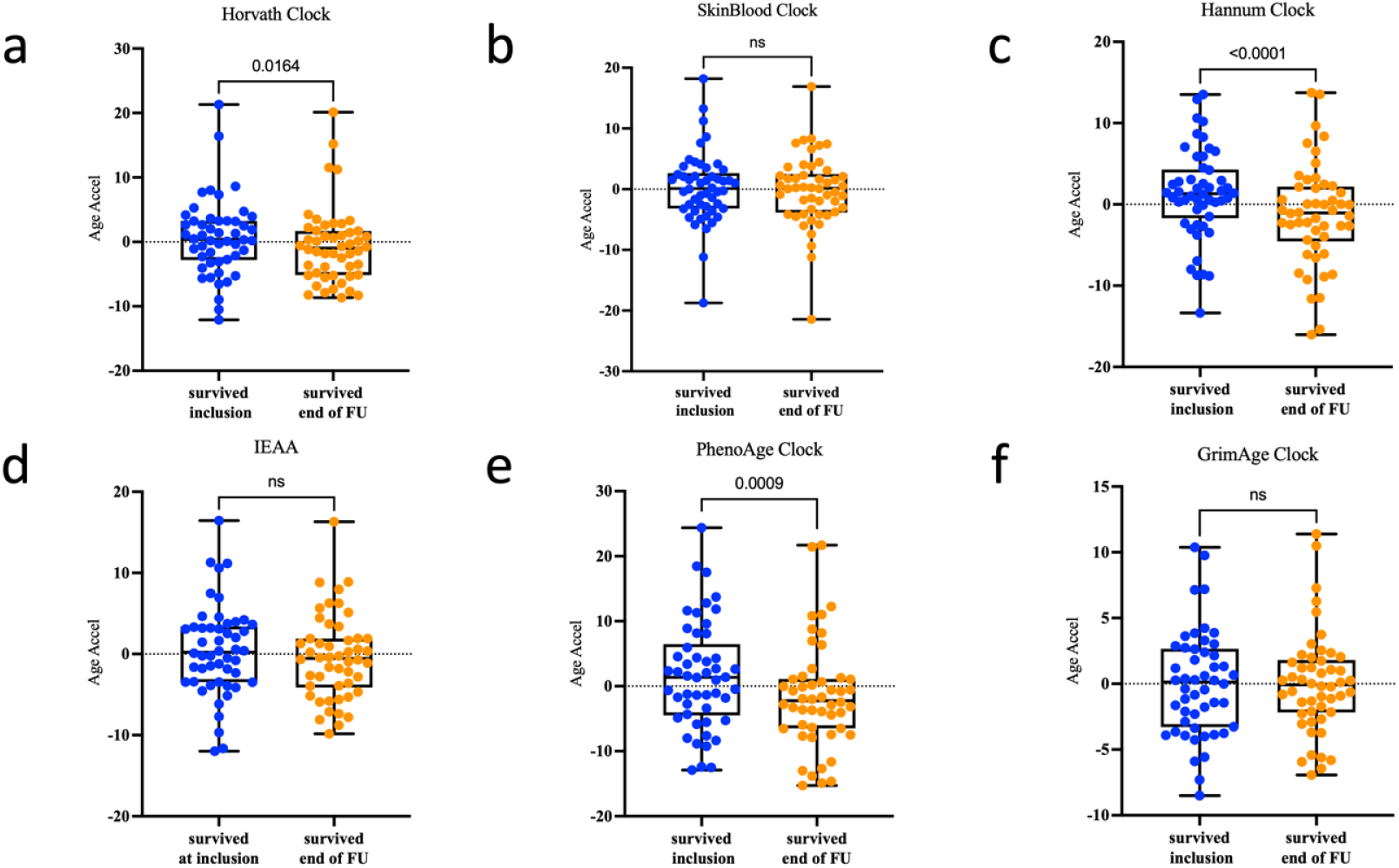
Epigenetic age acceleration at inclusion to the last recorded DNAmAge before recovery i.e. end of Follow Up (FU). DNAm age acceleration in six epigenetic clocks (a-f) in the peripheral blood from 50 COVID-19-T1 survived patients. The y-axis shows the epigenetic age acceleration. The p-value is shown above the corresponding line. In the box plots, the lower and upper hinges indicate the 25th and 75th percentiles and the black line within the box marks the median value. Ns: non-significant.

**Fig. 3.**
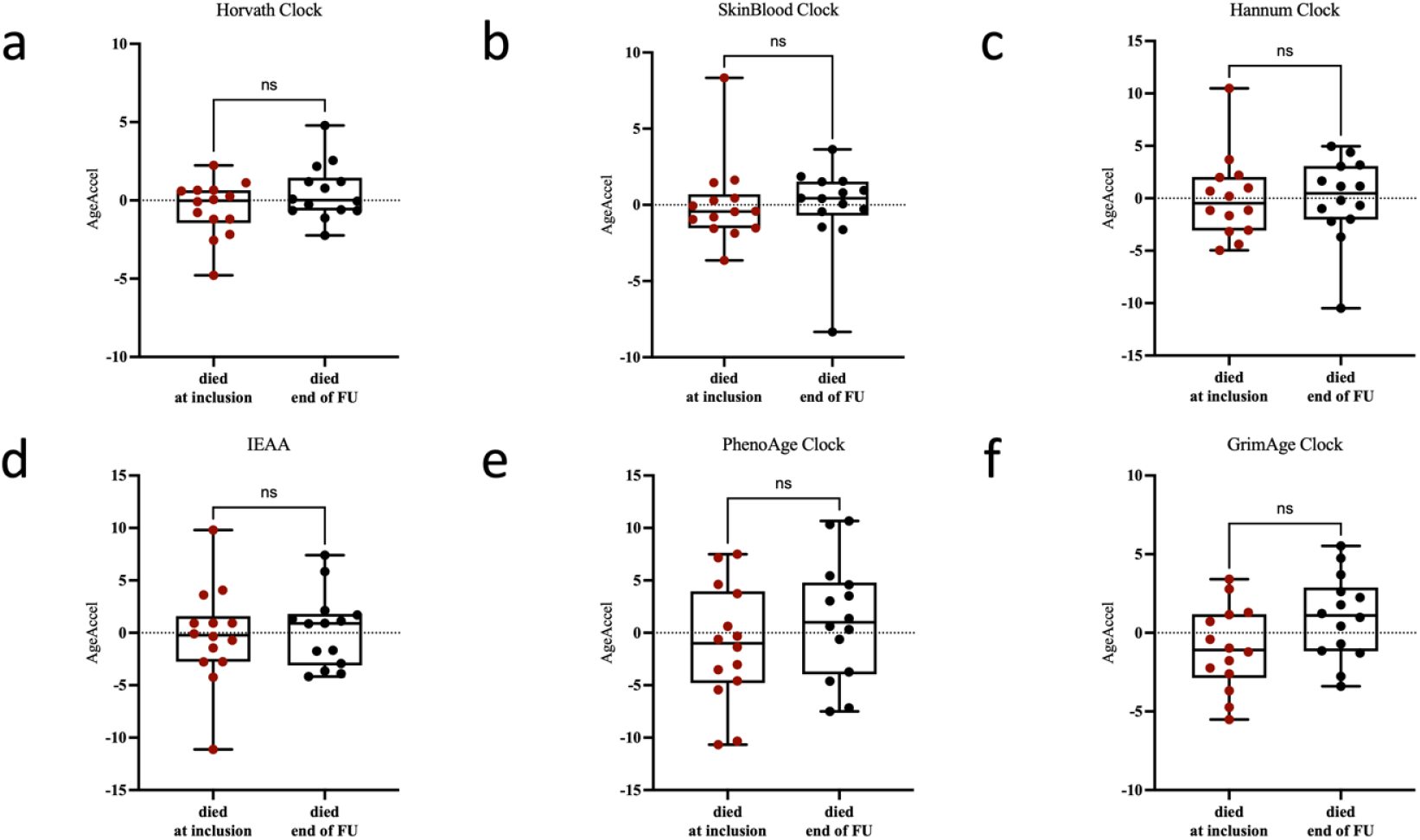
Epigenetic age acceleration in COVID-19 patients who died at inclusion vs the end of follow-up (FU) timepoint. Distribution of DNAm age acceleration in six epigenetic clocks (a-f) in the peripheral blood from 14 COVID-19 patients at inclusion vs end of follow up. The y-axis shows the epigenetic age acceleration. The p-value is shown above the corresponding line. In the box plots, the lower and upper hinges indicate the 25th and 75th percentiles and the black line within the box marks the median. Ns: non-significant.

**Fig. 4.**
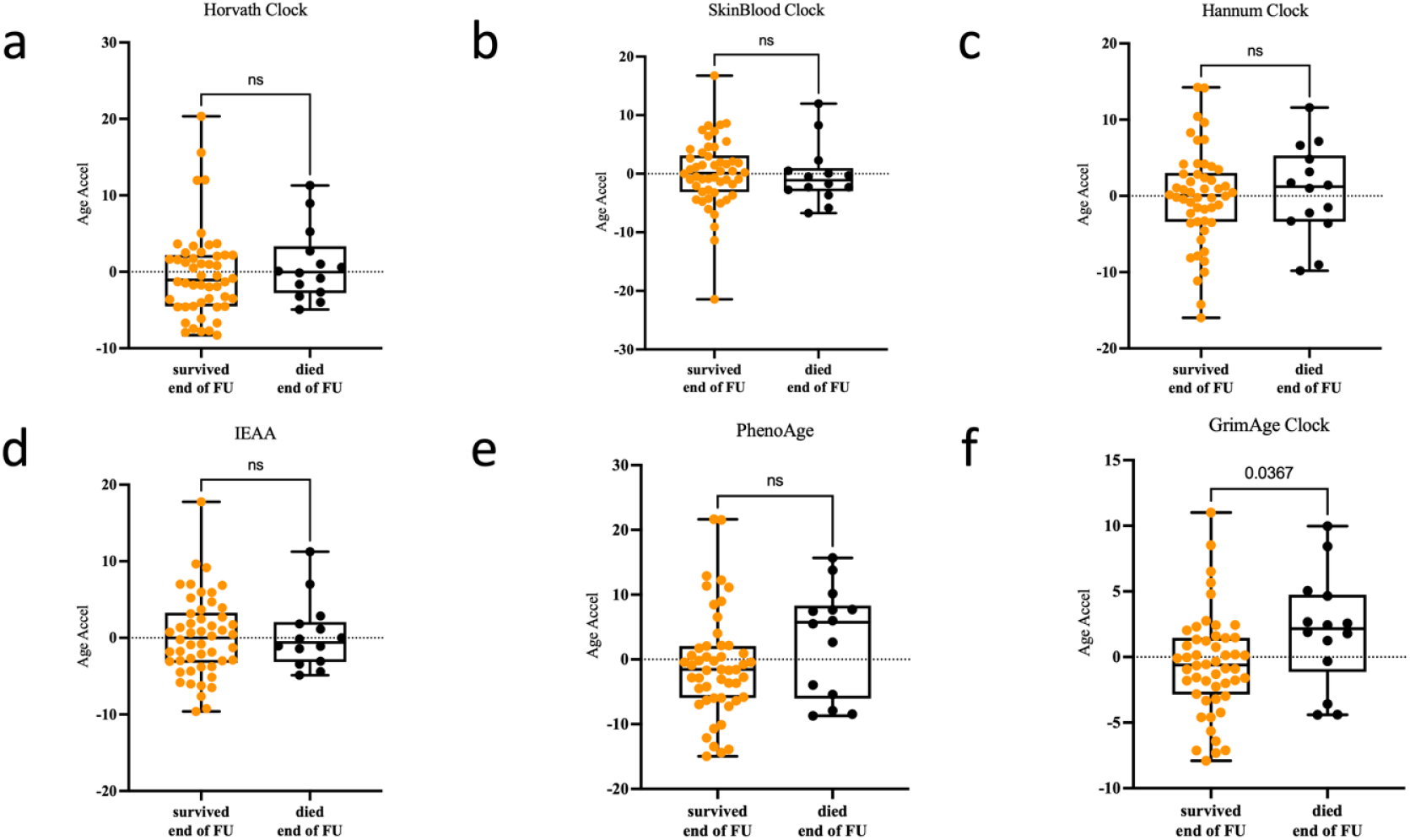
Epigenetic age acceleration in survived vs deceased COVID-19 patients at the end of follow up (FU) timepoint. Boxplots of DNAm age acceleration in six epigenetic clocks (a-f) in the peripheral blood from 50 surviving vs 14 deceased COVID-19 patients at the end of follow-up. The y-axis denotes the epigenetic age acceleration. P-value is shown above the corresponding line. In the box plots, the lower and upper hinges indicate the 25th and 75th percentiles and the black line within the box represents the median. Ns: non-significant.

### Longitudinal epigenetic aging acceleration across disease phases

Here, we used a linear mixed model to examine the dynamic acceleration of epigenetic age over continuous time points (T1-T5) of 50 COVID-19 patients who survived and 14 who died. Our analysis of dynamic EAA across these different time points showed no significant difference between COVID-19 patients who survived vs those who died following hospitalization in all of the employed epigenetic clocks. We additionally compared dynamic changes in EAA by calculating the difference between the epigenetic age acceleration at the end of follow-up and at inclusion time points in both groups, which we defined as “Change in AgeAcceleration”. In this analysis, we could observe a significant increase in EAA using both the Horvath and the PhenoAge clocks (p=0.0415 and 0.0207, respectively) (Fig. 5).

**Fig. 5.**
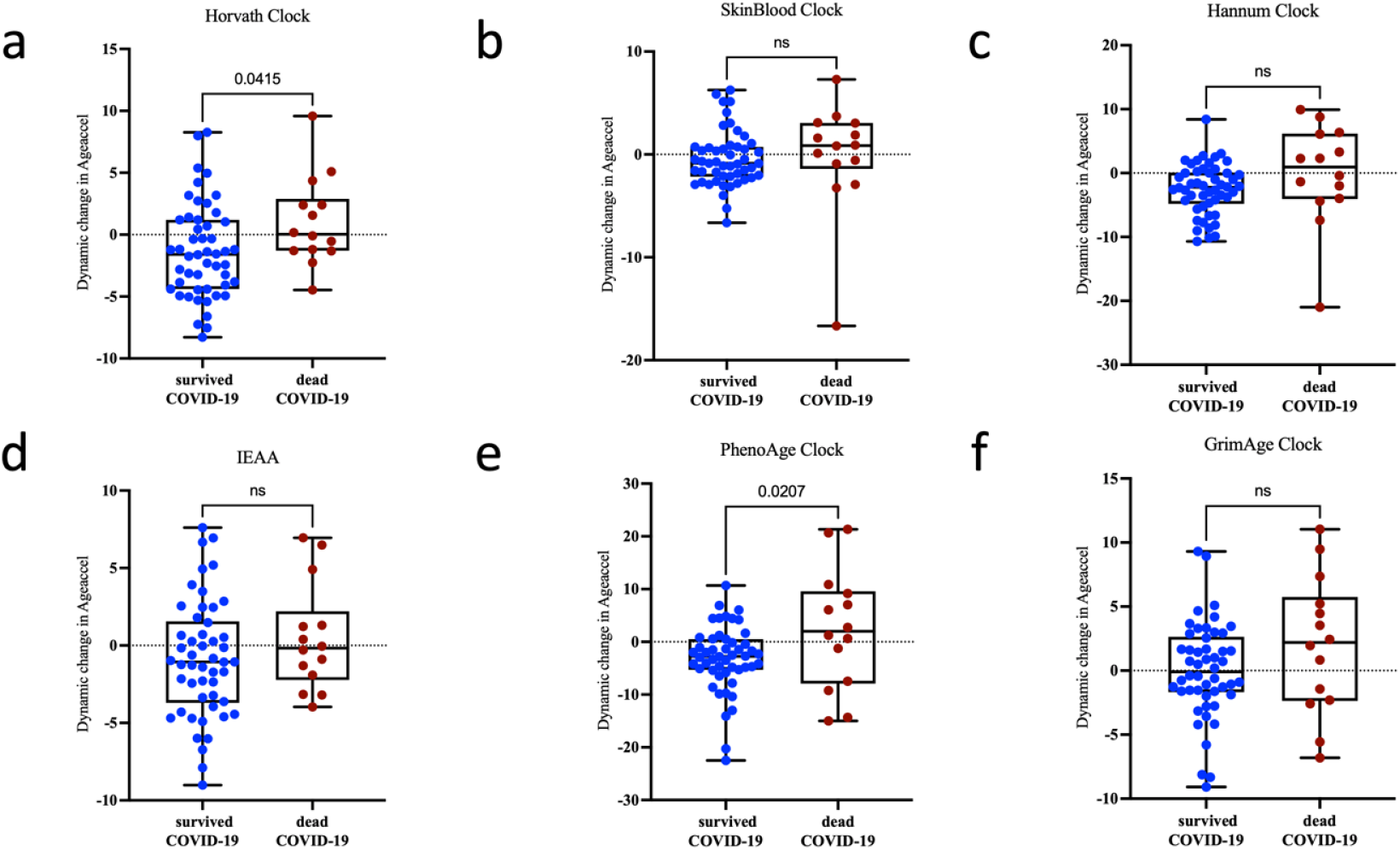
Dynamic change in epigenetic age acceleration in survived vs deceased COVID-19 patients between inclusion and the end of follow-up (FU) timepoint. Dynamic change in DNAm age acceleration in six epigenetic clocks (a-f) in the peripheral blood from 50 surviving vs 14 deceased COVID-19 patients. The y-axis denotes the difference in epigenetic age acceleration between the end of follow-up and inclusion (EAA end of follow-up – EAA inclusion). P-value is shown above the corresponding line. In the box plots, the lower and upper hinges indicate the 25th and 75th percentiles and the black line within the box represents the median. Ns: non-significant.

### Telomere attrition in COVID-19 patients

We calculated telomere length using the surrogate marker DNAm TL to compare accelerated telomere attrition in the studied cohort. The initial analysis of COVID-19 samples vs controls revealed no significant difference in DNAmTL attrition (Fig 6a). When comparing EAA between inclusion and end of follow-up, we observed no changes in DNAmTL attrition in the recovered critically ill COVID-19 patients. However, we detected a significant DNAmTL attrition in the deceased COVID-19 patients at end-of-follow-up (p= 0.0015) indicating a measurable reduction in telomere length in these individuals (Fig 6b-e). Finally, we compared dynamic differences in telomere attrition by calculating the variable “Dynamic Change in TL attrition acceleration” after subtracting TL attrition acceleration at inclusion from the end of the follow-up. Comparing Dynamic Change in TL attrition acceleration showed a significant reduction in TL attrition acceleration in the deceased COVID-19 patients between the two time-points (p=0.0077; Fig. 6f). Nevertheless, when we applied a mixed linear model over the continuous time points (T1-T5) for TL attrition acceleration, we did not observe any difference between the deceased vs surviving COVID19 patients.

**Fig. 6.**
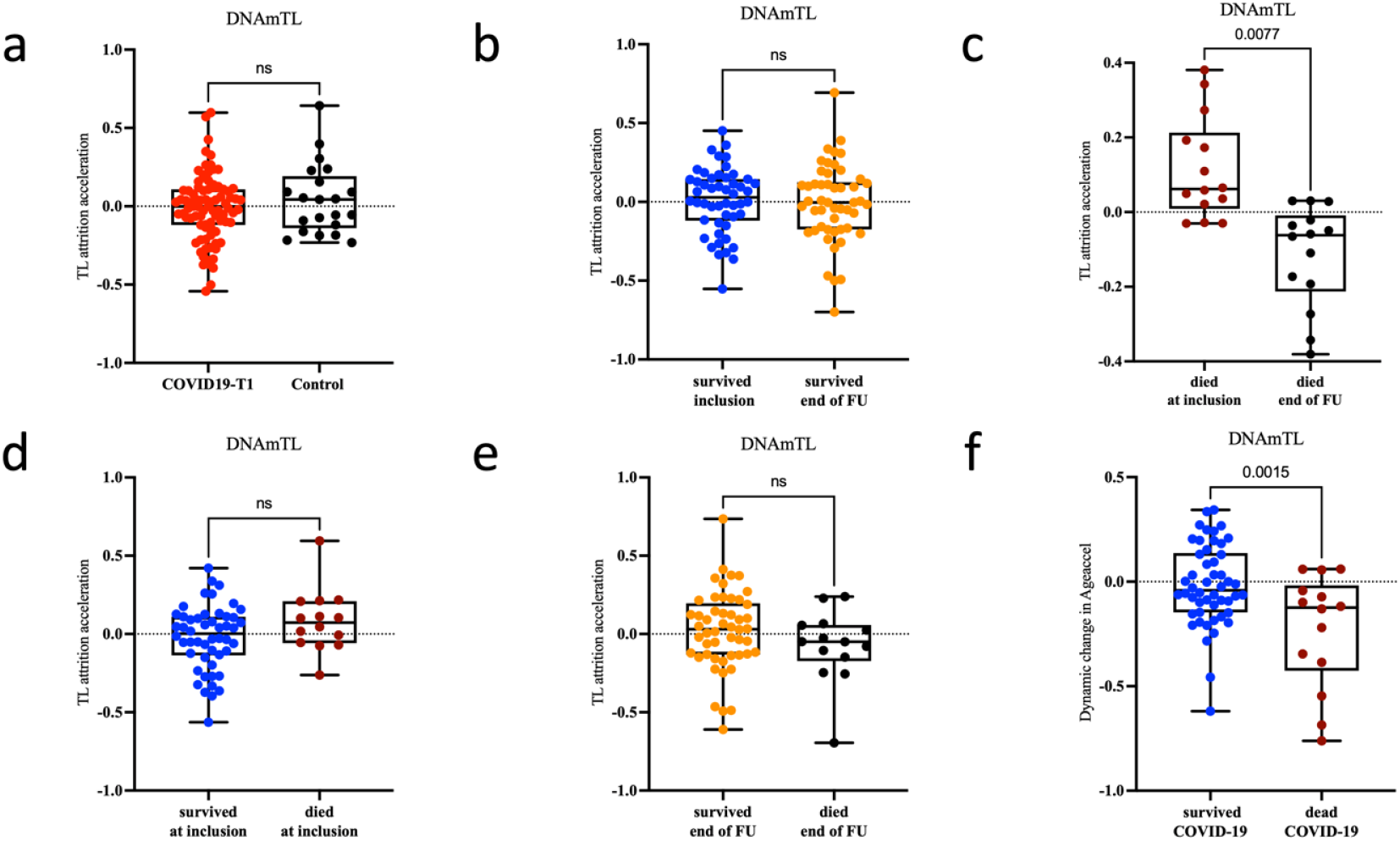
Telomere attrition in Covid-19 patients. Telomere attrition acceleration measured via the DNAm TL surrogate marker (a-f) in the peripheral blood of COVID-19 patients. Telomere attrition acceleration in **a)** COVID-19 patients at baseline (T1) vs controls **b)** in surviving COVID19 patients at inclusion vs the last recorded measurement at end of Follow Up (FU) **c)** in deceased COVID-19 patients at inclusion vs end of FU **d)** in survived vs deceased patients at T1and **e)** at the end of FU. **f)** Dynamic change in telomere attrition acceleration between the end of follow-up and inclusion. The y-axis denotes telomere attrition acceleration apart panel (f), representing a change in telomere attrition acceleration between the end of FU and inclusion. P-value is shown above the corresponding line. In the box plots, the lower and upper hinges indicate the 25th and 75th percentiles and the black line within the box represents the median. Ns: non-significant.

## Discussion

In this longitudinal study, we performed an epigenetic age acceleration analysis in critically ill COVID-19 patients with ARDS receiving mechanical ventilation. In addition, we studied telomere length in the same cohort using the surrogate marker DNAmTL. Overall, we observed increased epigenetic age acceleration in critically ill COVID-19 patients compared to non-infected controls. This EAA was detected using one of the first-generation clocks (Hannum clock) as well as the second-generation clocks (PhenoAge and GrimAge) (18, 19, 22). Second-generation clocks were trained to predict surrogate measures of phenotypic age instead of only chronological age and are therefore more accurate in predicting biological aging, morbidity, and mortality (10). Our findings reveal that individuals with increased EAA are more predisposed to develop severe COVID-19 symptoms including ARDS. A handful of studies reported an effect of viral infection including HIV and SARS-Cov2 on epigenetic aging (14, 23, 24). Corley et al. first reported increased DNAm age measured via the second-generation GrimAge clock in severe COVID-19 cases. In contrast, Franzen et al did not observe EAA in hospitalized COVID-19 patients with and without ARDS using four first-generation DNAm age predictors (25, 26). More recently, Cao et al. reported accelerated epigenetic aging associated with SARS-CoV-2 infection and COVID-19 disease severity. The study used multiple epigenetic clocks including first- and second-generation clocks and looked at a large cohort of 194 patients with mild/moderate symptoms and 213 severe COVID-19 patients (21). Their findings align with our study’s observations, particularly for certain epigenetic clocks.

Furthermore, we could observe that reversing the epigenetic age acceleration (i.e. epigenetic deceleration) measured by the Horvath, Hannum, and PhenoAge clock is associated with recovery in severe COVID-19 patients following hospitalization. In contrast, patients who died after ICU admission did not show any differences in epigenetic aging at inclusion to the last recorded DNAm age before death. This indicates that COVID-19 alters epigenetic aging and decelerating this EAA would lead to recovery following ICU admission. Recently, Poganik et al. reported transient changes in epigenetic aging in patients with severe COVID-19 and during surgery, where patients exhibited an increase in biological aging following exposure to stress that was later reversed after recovery (27). This aligns with the findings from our study where EAA was reversed in patients that recovered following hospitalization. Similarly, the previously mentioned study by Cao et al. analyzed dynamic changes in EAA during multiple disease phases, where the authors observed DNAm age acceleration at the initial phase of infection to be partly reversed at the later convalescence phase (21). One important question is whether the observed EAA is causally related to disease risk and severity or is a consequence of SARS-CoV-2 infection. In this context, a single Mendelian randomization study looked at the causal relationship between COVID-19 and epigenetic aging. However no causal association was observed between epigenetic age and COVID-19 susceptibility (28).

Interestingly, in our study we could observe that COVID-19 patients that died following mechanical ventilation exhibited significant telomere shortening at the last recorded timepoint before death when compared to the first timepoint at ICU admission. This suggests that critically ill patients with telomere attrition are more likely to die during hospitalization. Similarly, dynamic change in TL attrition acceleration between end of follow-up and inclusion revealed significant differences between survived vs deceased patients. Despite conflicting data, several studies including meta-analyses showed increased risk for all-cause mortality to be associated with shorter telomeres in the general population (29-31). It is important to mention that we could not identify differences in telomere attrition between COVID-19 patients and controls, which is in contrary to other studies reporting telomere shortening following SARS-CoV-2 infection (21, 32). Similar to our study, Cao et al. used the DNAmTL surrogate marker to measure telomere length (21), whereas Mongelli et al. employed a qPCR-based assay for absolute telomere length quantification (32). The output of DNAmTL is known to exhibit a moderately strong association with telomere length measured using qPCR or southern blotting in the blood (20). However, our findings align with the study of Franzen *et al*. where lymphocytes from severe COVID-19 patients did not show a significant acceleration of telomere attrition (26).

Regarding dynamic changes in EAA during disease phases, the study by Cao et al. looked at a previously published dataset of only six COVID-19 cases and six uninfected controls (21, 33) were analyzed. To our knowledge, our study is the first to look at EAA following a large-scale longitudinal follow-up of severe COVID-19 cases with ARDS and to determine the relationship between EAA/Telomere attrition and outcome (survival or death). Nevertheless, one of the main limitations of this study is the lower number of non-Covid controls (N=21) and the limited number of studied samples who died following ICU admission. In addition, we could only determine an association between accelerated epigenetic aging and COVID-19, however understanding the causal relationship between them was not within the scope of the current study.

In conclusion, we demonstrate that severe COVID-19 is associated with a significant increase in DNA methylation age but not in DNAm telomere attrition. In addition, we could also detect an association between EAA and recovery/death following hospitalization in COVID-19 patients. Similarly, an association between DNAm telomere attrition with outcome was also observed. Future studies will need to explore whether epigenetic age acceleration is causally linked to disease severity and outcome following hospitalization, and the underlying biological mechanisms behind this association. Furthermore, understanding the long term effects and consequences of accelerated aging in recovered COVID-19 patients is essential and urgently needed.

## Methods

### Ethical Approval

The study is part of the “Immune Profiling of COVID-19 patients Admitted to ICU study (IMPROVISE) (clinicaltrial.gov identifier NCT0447313). The Institutional review board at Hamad Medical Corporation (HMC) and Weill-Cornell approved the study with record numbers MRC-05-007 and 20-00012. All research was conducted in accordance with the ethical principles of the Declaration of Helsinki. All participants enrolled in this study or their legal guardians signed a consent form.

### Participants

In total, 100 critically ill COVID-19 patients with acute respiratory distress syndrome (ARDS) receiving mechanical ventilation in the intensive care unit (WHO clinical progression scale 7-9) (34) and 24 non-COVID controls from HMC blood donor unit were selected. The ethnicity of the patients and controls were matched. Our previous study by Bradic et al. provides detailed inclusion and exclusion criteria for the enrolled individuals (35). Following their admission to the intensive care unit (Time point 1: T1), COVID-19 patients were monitored at four different time points, including days 7 (T2), 14 (T3), 21 (T4), and 60 (post-T4). The follow-up was carried out in accordance with the guidelines of the WHO Working Group on the Common Outcome Measure Set for the COVID-19 Clinical Research (34). Blood samples were collected at each time point unless the patient has died or recovered at the end of the follow-up. Recovery was defined as per WHO clinical criteria of less or equal to 5, discontinuing mechanical ventilation, and discharge from the ICU (34).

### Methylation array data processing

Samples were processed on the Illumina Infinium MethylationEPIC Beadchip (EPIC array) which interrogates > 850,000 CpG sites across the human genome including extensive coverage of genes, promoters, CpG Islands, and enhancers. Raw IDAT files for a total of 288 samples were obtained and processed further (35). The RnBeads package (36) was used for quality control and noob data normalization was performed using the minfi package (37). Following data filtration, 12 samples were excluded from further analysis since they failed to meet quality criteria. After data normalization, the methylation β value for each CpG site was extracted and used as input to calculate the epigenetic age using various clocks.

### Epigenetic age calculation and DNAmTL estimation

DNAm age was estimated using various epigenetic clocks such as the Horvath pan-tissue (17), PhenoAge (18), GrimAge (19), and SkinandBlood (38) clocks via the web-based epigenetic clock calculator (https://dnamage.clockfoundation.org/). In addition, intrinsic epigenetic age acceleration (IEAA) was measured since it reflects epigenetic age independently of age-related changes in blood composition (39). Epigenetic age acceleration (EAA) is defined as the deviation between epigenetic age and chronological age. EAA was calculated based on the residuals from regressing DNAm age on chronological age after correcting for BMI and gender. Furthermore, DNAmTL (20) was estimated where the deviation between DNAm TL and chronological age is defined as DNAm TL attrition acceleration. This measurement was calculated by adjusting for BMI and gender as covariates. All statistical analyses were performed using RStudio version (2023.3.1) and Prism Software (version 9.51).

### Statistical analysis

The correlation between DNAm age, DNAm TL, and chronological age of the samples was evaluated using Pearson correlation (*R*). To compare samples within the same group (individuals who survived COVID-19 or individuals who died from COVID-19) at two different time points (T1 and end of follow-up), a paired t-test was conducted. Additionally, an unpaired t-test was employed to compare COVID-19 samples to control samples, and to compare individuals who survived versus those who died at the inclusion or at the end of the follow-up period. A dynamic age acceleration linear mixed model was conducted to assess changes in epigenetic age in relation to outcome (survival or death). Subsequently, statistical significance was evaluated using a chi-square test. All statistical analyses were performed using RStudio version(2023.3.1) and Prism Software (version 9.51). P values < 0.05 were considered statistically significant.

## Availability of data and materials

The datasets used in the current study are available from the corresponding authors upon request.

## Contributions

Y.B., S.T., and M.B. performed experiments. Y.B. and F.H.A. performed bioinformatic analyses. M.S. supervised statistical analysis performed by Y.B.. A.A.H. and C.A.K. collected study samples. Y.B., A.M., and N.E.H wrote the manuscript. M.S., A.M., C.A.K, and N.E.H critically reviewed and edited the manuscript. N.E.H designed the study.

## Ethics declarations

### Ethics approval and consent to participate

The study is part of the “Immune Profiling of COVID-19 patients Admitted to ICU study (IMPROVISE) (clinicaltrial.gov identifier NCT0447313). The study was approved by the Institutional review board at HMC and Cornell with record number MRC-05-007 and 20-00012. All research was conducted in accordance with the ethical principles of the Declaration of Helsinki. All participants enrolled in their study or their legal guardians signed a consent form.

### Consent for publication

Not applicable

## Supplementary Information

**Supplementary Figure 1.**
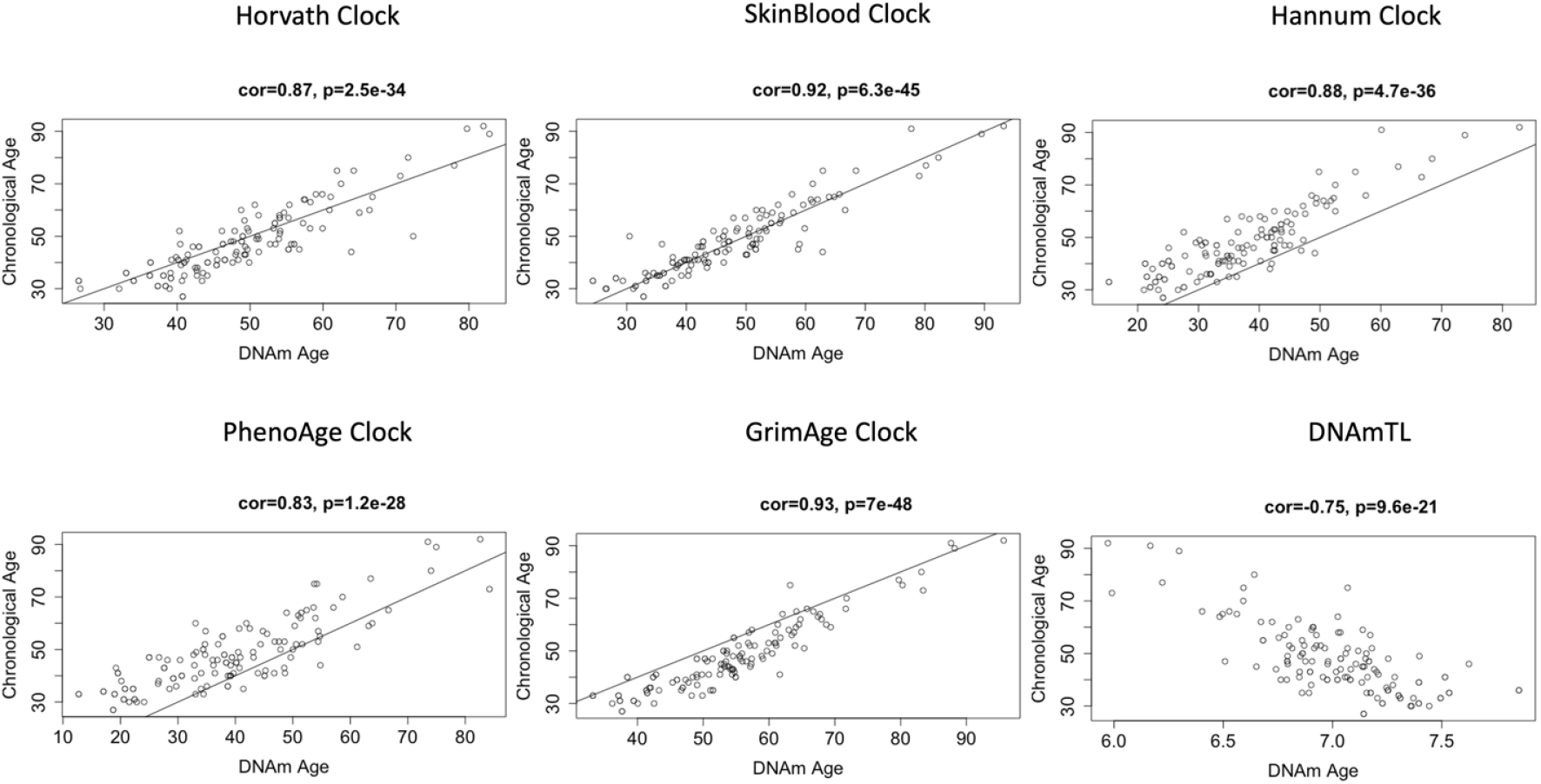
Correlation of chronological age with DNA methylation age using Horvath, Hannum, SkinandBlood, PhenoAge, and GrimAge clocks and the DNA methylation-based telomere length (TL) estimator.

